# Single-Platform Nanopore Sequencing Enables Diploid Telomere-to-Telomere Genome Assembly and Haplotype-Resolved 3D Chromatin Maps

**DOI:** 10.64898/2026.03.19.712851

**Authors:** Caspar Gross, Ramya Potabattula, Fubo Cheng, Sarah Leuchtenberg, Hanna Sophie Hartung, Beate Kristmann, Elena Buena-Atienza, Nicolas Casadei, Stephan Ossowski, Olaf Riess

## Abstract

Telomere-to-telomere (T2T) genome assembly has transformed human genomics by resolving centromeres, segmental duplications, and other previously inaccessible regions. However, most diploid T2T assemblies rely on the combination of multi-platform sequencing strategies including short read genome sequencing, PacBio HiFi, Oxford Nanopore ultra-long reads, and chromatin conformation capture data (Hi-C), limiting both scalability and accessibility. Here, we present a streamlined Nanopore-only workflow for diploid human T2T assembly using three ultra-long and one Pore-C PromethION flow cell per individual. Across 23 genetically diverse individuals, we generated 360 gapless chromosomes and 446 near-complete T2T scaffolds, achieving median consensus accuracy of QV50 without Duplex sequencing or hybrid polishing. Assembly continuity, gene completeness, and structural variant detection were comparable to multi-platform Human Pangenome Reference Consortium assemblies. Pore-C data enabled chromosome-scale haplotype phasing without parental information and supported generation of haplotype-resolved chromatin contact maps. Integrated methylation and 3D genome analyses revealed allele-specific chromatin organization at imprinted loci and clear signatures of X-chromosome inactivation. Our openly accessible dataset expands public T2T resources and demonstrates that reference-grade diploid assemblies, phased methylomes, and 3D genome maps can be derived from a single sequencing platform. This approach reduces technical barriers and supports scalable population and functional genomics in the T2T era.

## Introduction

The recent completion of a telomere-to-telomere (T2T) human reference genome^1^ and the subsequent assembly of the first fully phased diploid human genome^2^ marked a milestone in genomics, resolving previously inaccessible regions including centromeres, segmental duplications, and large repetitive arrays. This achievement lays the foundation for the Human Pangenome Reference Consortium (HPRC), which aims to construct a comprehensive human pangenome based exclusively on high-quality T2T assemblies. The initial pangenome draft^3^ has already demonstrated the value of multiple complete assemblies for resolving structural variation, characterizing centromere architecture^4^, and improving variant discovery in complex genomic regions^5^. Population-specific T2T assemblies further highlight the importance of complete references for understanding genomic diversity across a wide range of ancestries^6,7^.

Current strategies for generating diploid T2T assemblies typically rely on the integration of multiple sequencing platforms using assembly methods such as *Verkko*^8^ or *hifiasm*^9^. High-accuracy long reads, most commonly PacBio HiFi, are used to construct the primary assembly graph, while Oxford Nanopore Technologies (ONT) ultra-long reads provide spanning information to resolve repeats and close remaining gaps. Additional datasets such as Hi-C^10^, Pore-C, Strand-seq^11^, or parental trio sequencing are then employed for haplotype phasing and scaffolding. While highly effective, these multi-platform approaches are cost intense and come with increased-laboratory complexity and logistical requirements, thus limiting broad accessibility outside large genome centers. Recently, Nanopore-only strategies have been explored as a potential alternative. Near-complete assemblies have been generated without ultra-long reads^12^, and gapless assemblies have been demonstrated using combinations of ultra-long reads, Pore-C, and Nanopore Duplex sequencing^13^. Duplex reads achieve improved per-read accuracy by combining signals from both DNA strands; however, their yield remains limited and the workflow adds additional complexity and cost. Thus, whether high-quality diploid T2T assemblies can be generated using only standard ultra-long Nanopore reads—without HiFi data and without reliance on Duplex reads—remains an open question.

Beyond linear genome reconstruction, comprehensive diploid assemblies provide an opportunity to integrate higher-order genome organization and epigenetic information. The three-dimensional (3D) chromatin architecture of the genome is organized into topologically associating domains (TADs)^14^, which constrain regulatory interactions and are frequently demarcated by CTCF binding sites and cohesin-mediated loop extrusion^15,16^. While chromatin structure is dynamically re-established during the cell cycle, allele-specific TAD configurations can arise through epigenetic mechanisms such as imprinting-mediated differential methylation at CTCF binding sites^17^. Chromatin conformation capture (3C) technologies, particularly Hi-C^10^, have enabled genome-wide interrogation of chromosomal contacts and are routinely used for scaffolding and phasing in de novo assemblies. Pore-C extends this principle by coupling proximity ligation with long-read Nanopore sequencing, thereby capturing multi-way contacts within single concatemers^18^. Compared to short-read Hi-C, Pore-C enables higher-resolution mapping of mid-range chromatin interactions and provides improved long-range scaffolding power. When combined with phased heterozygous variants, Pore-C data allow assignment of contacts to individual haplotypes^19^, enabling haplotype-resolved maps of 3D genome organization^20^. Importantly, Nanopore sequencing simultaneously preserves native DNA methylation information^21^, allowing joint reconstruction of phased genome sequence, chromatin topology, and methylome from a single technological platform.

Here, we demonstrate that high-quality, near-complete diploid human T2T assemblies can be generated using only Nanopore sequencing on four PromethION flow cells, without PacBio HiFi data and without Duplex reads. Using three flow cells of ultra-long R10.4 reads and one flow cell of Pore-C data, we assembled 31 of 46 chromosomes gaplessly (Sample T2T17) and achieved assembly quality metrics comparable to hybrid Nanopore–HiFi approaches. In parallel, we generated haplotype-phased maps of chromatin architecture and DNA methylation. Our streamlined workflow substantially reduces cost, laboratory complexity, and hands-on time while delivering integrated structural and functional genome information, providing a scalable framework for population-scale T2T genomics.

## Results

### A streamlined Nanopore-only workflow enables diploid T2T assembly from four flow cells

We generated diploid telomere-to-telomere (T2T) assemblies for 23 individuals, including two parent-child trios of German ancestry, using a Nanopore-only workflow (Fig. 1). Our standard operating procedure for T2T assembly uses three PromethION R10.4.1 flow cells of ultra-long (UL) reads and one to two flow cells of Pore-C data (>50Gb yield), resulting in a total of four flow cells per genome under optimal conditions (Supplementary Fig. 1). No PacBio HiFi, Nanopore Duplex-reads, Strand-seq, or parental data were required for routine assembly.

**Figure 1:**
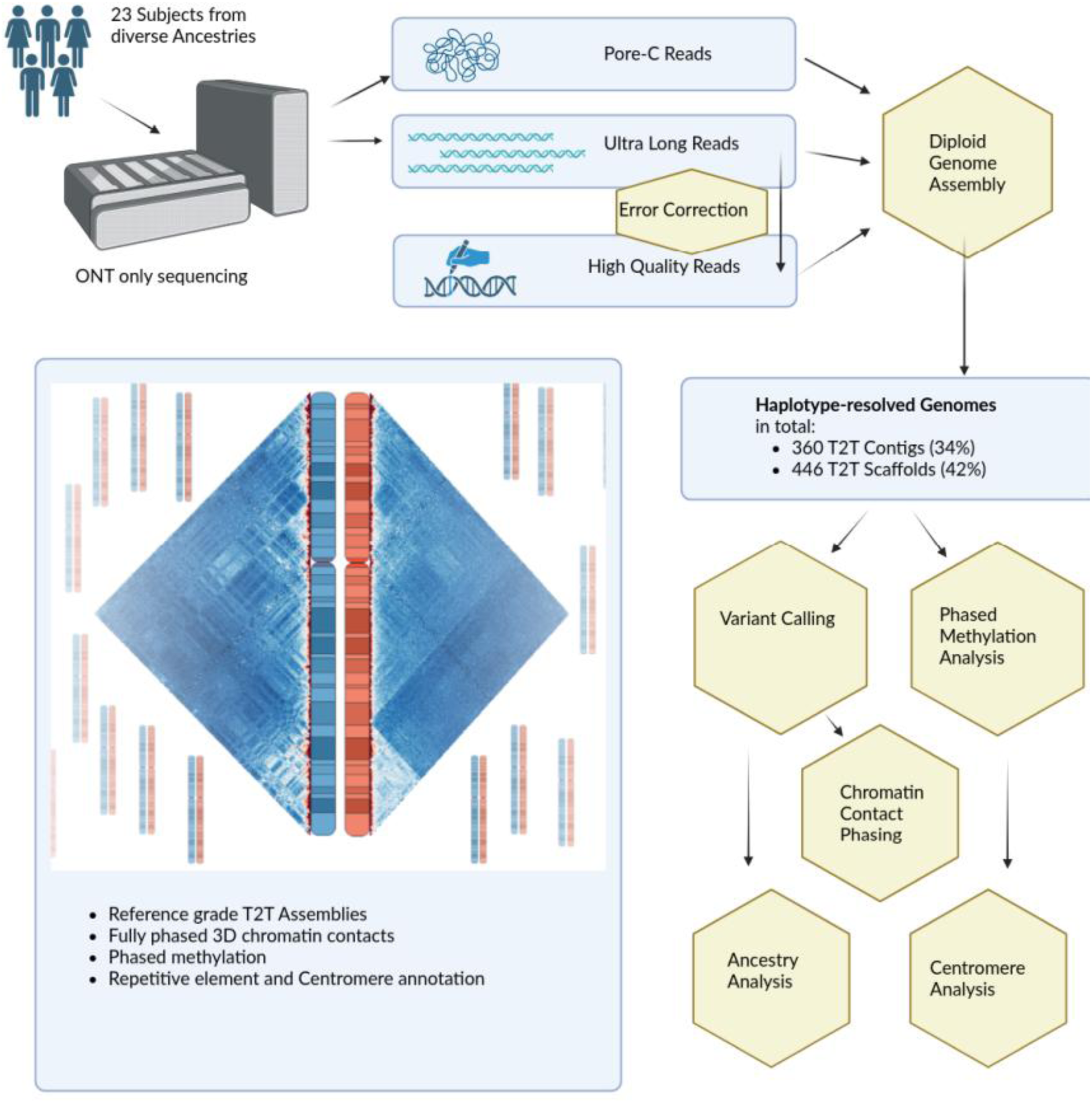
Graphical abstract. Graphical overview of the experimental and computational workflow. Fresh peripheral blood is processed to generate ultra-long (UL) Nanopore libraries and Pore-C libraries. UL reads are error-corrected to produce high-quality (HQ) reads for graph-based de novo assembly using Verkko. Pore-C data enables chromosome-scale scaffolding and haplotype phasing. Downstream analyses include assembly quality control, phased variant detection, centromere characterization, haplotype-resolved methylation profiling, and 3D chromatin contact map generation. Our standardized T2T-3D procedure integrates all steps into a unified workflow.

To ensure optimal long-read performance, we refined UL library preparation to maximize reads exceeding 100 kb while maintaining high total yield (Supplementary Fig. 2). UL reads were bioinformatically error-corrected using *HERRO* to generate high-quality (HQ) reads, which were used as substitutes for HiFi reads in *Verkko* assembly. Reads were filtered to retain 50× HQ coverage and 70× UL coverage, prioritizing read length and retaining only reads longer than 80kb to maximize repeat resolution (Supplementary Fig. 3). Pore-C data were preprocessed using ONT’s *wf-pore-C* pipeline.

All data processing steps were automated combining preprocessing, assembly, phasing, quality control, variant calling, methylome and 3D genome analysis. Assemblies were generated using *Verkko v2.2.1* with Pore-C–based or Trio-based phasing. Furthermore, we generated haplotype-phased assemblies using the *hifiasm* method for benchmarking purposes. For each assembler and phasing combination, we generated a wide range of quality measures to demonstrate the performance of Nanopore-only compared to hybrid-platform T2T assemblies.

### High rates of gapless chromosome assembly across 23 diploid genomes

We classified *Verkko*-assembled chromosomes as T2T-scaffolds if they spanned ≥95% of the reference chromosome and contained telomeric repeats at both ends. Gap-free T2T-scaffolds were further classified as T2T contigs. The best-performing genome (T2T17) achieved 31 gapless T2T-contigs and 8 additional near-complete T2T-scaffolds (Fig. 2B). Across all 23 individuals, we reconstructed 360 gapless chromosomes (34% of all haplotypes) and an additional 446 near-complete T2T-scaffolds (42%). Many remaining gaps were small (<50 kb) and occurred as single isolated regions within otherwise continuous scaffolds (Supplementary Fig. 4).

**Figure 2:**
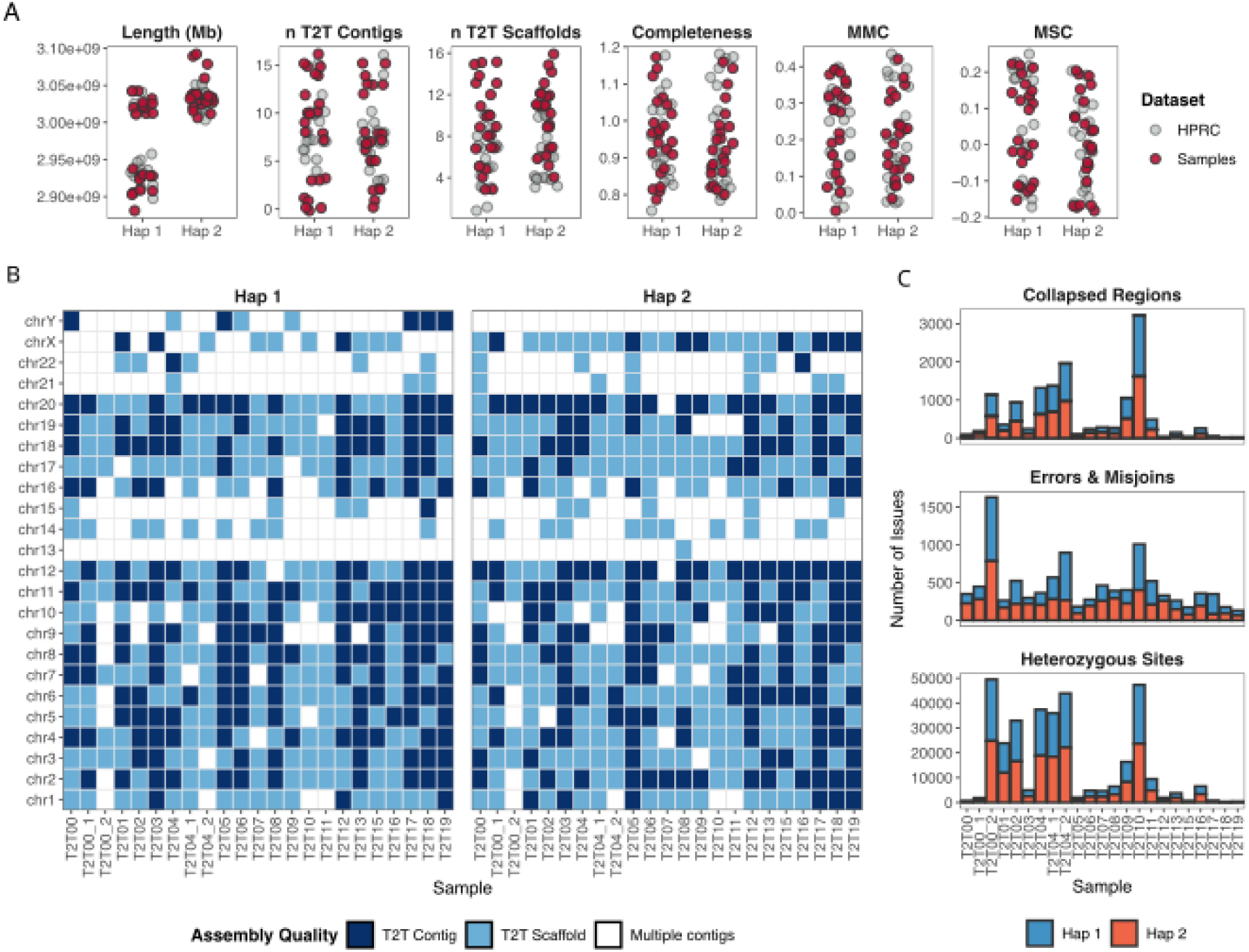
Nanopore-only assemblies achieve hybrid-grade T2T quality metrics. **(A)** Comparison of 23 Nanopore-only assemblies (red) with 20 multi-platform HPRC assemblies (gray) using identical quality control metrics. Shown are assembled genome length, number of gapless T2T contigs, number of near-complete T2T scaffolds, gene completeness, fraction of missing multi-copy genes (MMC), and fraction of missing single-copy genes (MSC). Distributions are comparable across datasets. **(B)** Chromosome-level assembly continuity across both haplotypes for all individuals. Dark blue indicates gapless T2T contigs; light blue indicates near-complete T2T scaffolds containing ≥1 gap; white indicates incomplete chromosomes. **(C)** Potential assembly issues detected by NucFlag following realignment of HQ reads to assembled genomes. Each bar represents one haplotype; variability reflects differences in sequencing and library preparation performance across samples.

Autosomal chromosomes 1–12 and 16–20 were consistently resolved to T2T contig or scaffold level across individuals. As expected, acrocentric chromosomes (13, 14, 15, 21, and 22) were more challenging; however, complete assemblies were achieved for specific acrocentrics in multiple individuals (e.g., chromosome 15 in T2T18; chromosome 22 in T2T04 and T2T16). In individual T2T18, all chromosomes except chromosome 13 were assembled to T2T scaffold or contig quality, including chromosome Y.

Performance improved over time as ultra-high molecular weight (UHMW) DNA and UL library preparation was optimized (Fig. 2C), with later samples achieving higher T2T-contig counts independent of total sequencing coverage (Supplementary Fig. 1). Instead, the proportion of reads exceeding 100 kb strongly correlated with assembly continuity (Supplementary Fig. 2).

### Nanopore-only assemblies achieve hybrid-grade reference-free quality metrics

We next evaluated reference-free assembly quality metrics using *paftools* and compared our Nanopore-only assemblies to multi-platform (PacBio + ONT) assemblies from HPRC. Across genome size, gene completeness, missing multi-copy genes (MMC), retained single-copy genes (MSC), and T2T-contig counts, our assemblies displayed distributions comparable to HPRC assemblies (Fig. 2A; Supplementary Fig. 5). Underperforming samples were primarily from early protocol iterations, supporting the importance of optimized UHMW DNA isolation.

To detect potential structural mis-assemblies, we re-aligned HQ reads and analyzed coverage and allele-frequency patterns using *NucFlag*^22^. Assemblies with high T2T-contig counts showed correspondingly low numbers of flagged regions (Fig. 2C). Independent validation using *Flagger* confirmed these findings (Supplementary Fig. 6), with flagged regions enriched in known difficult genomic loci (Supplementary Fig. 7).

### Base-level consensus accuracy reaches QV50 without Duplex reads

Consensus accuracy was evaluated using *Merqury* based on Illumina short-read k-mers, generated from re-alignment of >30x coverage per individual. For k=31, median QV across all samples was 51, corresponding to approximately 6 errors per megabase. Across k-mer sizes (21, 31, 37), median QV ranged from 48 to 60 (Supplementary Fig. 8A), demonstrating high base-level accuracy comparable to hybrid assemblies.

Trio-based phasing did not significantly improve QV. Notably, integration of ONT’s APK polishing kit reduced QV despite requiring additional sequencing, indicating that our streamlined workflow achieves optimal consensus without supplementary polishing steps (Supplementary Fig. 8B).

### Pore-C–based phasing matches trio phasing performance

Haplotype phasing was performed using Pore-C data for all samples. For two trios (T2T00, T2T04), we compared Pore-C–based phasing to trio-based phasing. Both approaches generated chromosome-scale haplotype blocks with consistent parental assignment across entire chromosomes (Supplementary Fig. 9). Using Pore-C, assignment of paternal or maternal origin is, as expected, arbitrary. Switch error rates and scaffold continuity were comparable between Pore-C and trio phasing using *Verkko*. In contrast, assemblies generated with *hifiasm* showed increased switch errors and reduced scaffold continuity relative to *Verkko*, despite producing slightly higher T2T-contig counts (Supplementary Fig. 9, 10, 11). These results demonstrate that Pore-C alone is sufficient for accurate haplotype phasing in Nanopore-only T2T assemblies.

### Diploid assemblies enable accurate phased variant discovery and ancestry inference

We aimed to confirm that the assembled genomes reflect the genetic variability expected from our cohort of diverse ancestries (Fig. 3A). We aligned each diploid assembly to T2T-CHM13 v2 to detect phased small and structural variants. For chromosomes assembled as T2T contigs or scaffolds, phase blocks spanned entire chromosomes.

**Figure 3:**
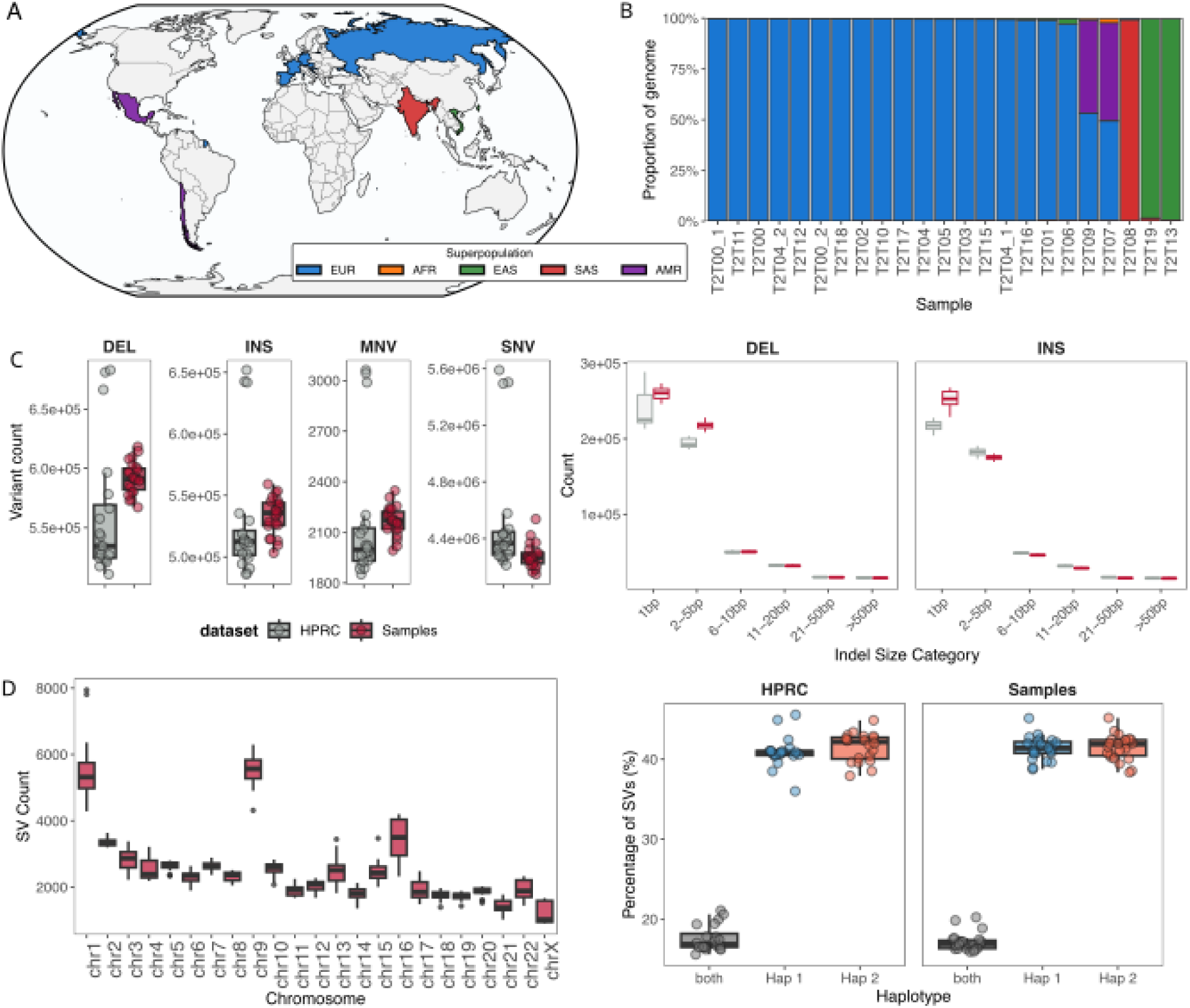
Phased variant discovery and ancestry validation in diploid T2T genomes. **(A)** Geographic origin of the 23 study participants, grouped by 1000 Genomes superpopulation (European [EUR], East Asian [EAS], South Asian [SAS], Admixed American [AMR]). **(B)** Ancestry inference of the assembled genomes, performed with RFMix using the 1000 Genomes Project reference panel. **(C)** Small variant counts in Nanopore-only assemblies (red) and 20 HPRC assemblies (gray), including single nucleotide variants (SNVs), insertions (INS), deletions (DEL), and multi-nucleotide variants (MNVs). Right panel shows size distributions of insertions and deletions. **(D)** Structural variant (SV) distribution across chromosomes, summed over all samples (left). Right panel shows allelic distribution of heterozygous and homozygous SVs in the study cohort compared to HPRC samples.

On average, we identified 4.4 million single nucleotide variants (SNVs) per individual. Transition/transversion ratios and heterozygosity rates were comparable to non-African HPRC genomes (Fig. 3C; Supplementary Fig. 12). A modest increase in small indels (1–5 bp) was observed, likely reflecting residual homopolymer-associated errors. Structural variant counts and chromosomal distributions closely matched HPRC samples (Fig. 3D), with ∼15% homozygous structural variants (SVs) and balanced heterozygous SV distribution across haplotypes.

Principal component analysis using a T2T-lifted 1000 Genomes reference panel confirmed accurate ancestry clustering consistent with self-reported origins (Supplementary Fig. 13).

Local ancestry inference of the assembled genomes, performed with RFMix using the coordinate lifted 1000 GP reference panel, confirmed the reported ancestries (Fig. 3B). Mixed ancestry was observed in only two samples with shared Admixed American (AMR) and European (EUR) ancestries, whereas all remaining samples match their expected 1000 Genomes superpopulation assignments.

### Haplotype-resolved Pore-C reveals allele-specific chromatin architecture

Using phased SNVs, we haplotagged Pore-C reads and generated haplotype-resolved contact maps. Approximately 50% of Pore-C reads could be assigned to a haplotype, yielding up to 1.6 billion phased contacts per sample (Fig. 4C). Additional Pore-C flow cells were sequenced for female samples T2T12, T2T13 and male samples T2T00 and T2T04 to improve the number of contacts at high resolutions. The sequencing statistics and quality metrics of Pore-C flow cells are presented in Supplementary Fig. 14 and Supplementary Fig. 15 respectively.

**Figure 4:**
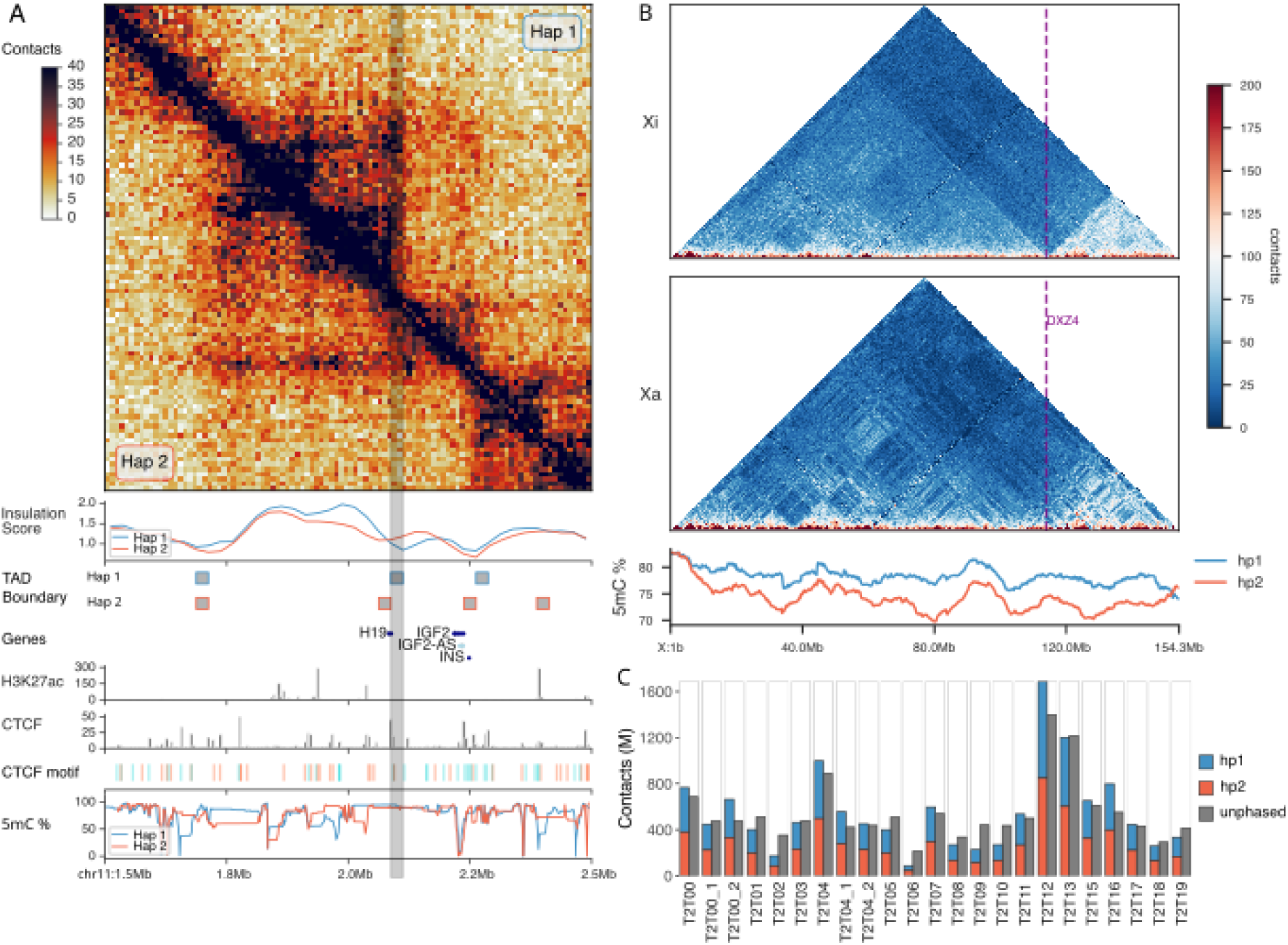
Haplotype-resolved Pore-C reveals allele-specific 3D chromatin organization. **(A)** Haplotype-specific chromatin contact maps (10 kb resolution) for the imprinted IGF2-H19 locus (chr11:1.5–2.5 Mb) in sample T2T12. Insulation scores highlight topologically associating domain (TAD) boundaries. Tracks include RefSeq gene annotation, H3K27ac signal, CTCF binding sites and motifs, and smoothed CpG methylation frequency (5mC). Distinct haplotype-specific chromatin structures are evident. **(B)** Chromosome-wide haplotype-separated contact maps (100 kb resolution) for chromosome X in T2T12, demonstrating differences between active and inactive X chromosomes. Bottom track shows smoothed CpG methylation frequency. **(C)** Number of chromatin contacts per sample. Unphased contacts were generated using wf-pore-c; haplotype-resolved contacts were obtained using a modified dip3d pipeline (cis interactions only).

Global chromatin architecture was highly concordant between haplotypes; however, allele-specific differences were detectable in specific loci. At the imprinted *IGF2-H19* locus, haplotype-specific TAD boundaries corresponded to differential methylation and CTCF binding (Fig. 4A). In samples of female participants, X-chromosome inactivation was evident as global differences in contact density and methylation between active and inactive X chromosomes (Fig. 4B). These results demonstrate simultaneous reconstruction of phased genome sequence, methylome, and 3D chromatin organization from a single sequencing platform.

### Nanopore-only assemblies reconstruct complete centromeric structures

Centromeres are notoriously hard to assemble correctly due to large clusters of alpha satellite (αSAT) tandem repeats^23^. We characterized centromeric regions using CenMap. The number of fully resolved centromeres per individual ranged from 2 to 42 (Fig. 5B), with highest resolution observed in assemblies with the greatest UL read length distribution.

**Figure 5:**
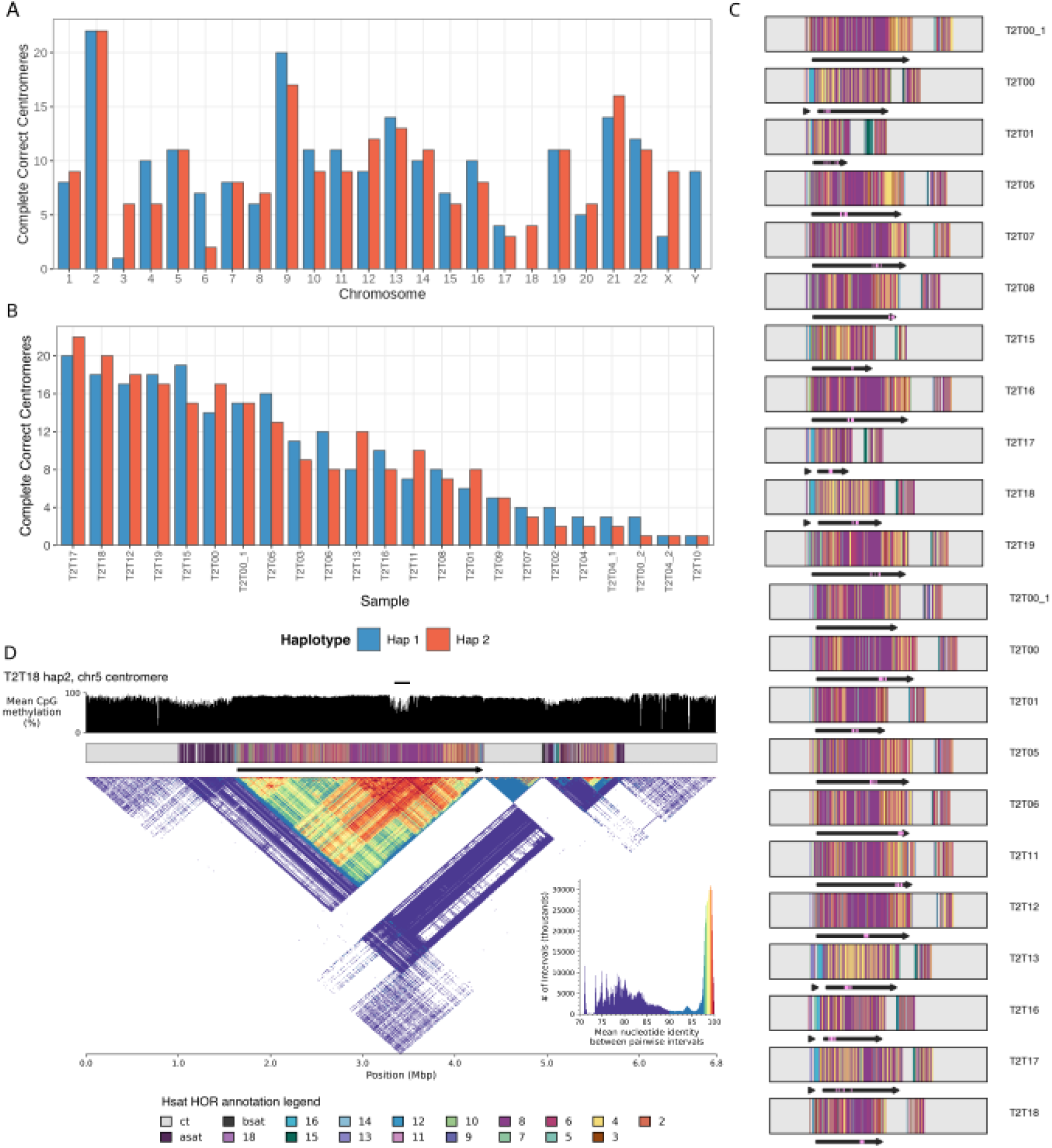
Nanopore-only assemblies reconstruct complete centromeric structures. **(A)** Number of fully resolved and structurally validated centromeres per chromosome across all 23 individuals and both haplotypes. **(B)** Total number of complete, validated centromeres per individual. **(C)** Example centromere structure for chromosome 5 across individuals. Colors denote higher-order repeat (HOR) classes. Black arrows indicate HOR arrays; centromeric dip regions (CDRs) are highlighted in pink. **(D)** Repeat homology plot for chromosome 5 centromere (haplotype 2, sample T2T18). Top track shows CpG methylation with CDR annotation. Second track shows HOR class annotation. Heatmap indicates pairwise nucleotide identity within satellite repeat arrays.

Consistent with prior studies, certain centromeres (e.g., chromosome 2) were reliably resolved, whereas others (e.g., chromosome 18) remained challenging (Fig. 5A). Extensive variation in centromere length and higher-order repeat (HOR) organization was observed between individuals (Fig. 5C), highlighting the rapid evolutionary dynamics in centromeric regions^24^.

The T2T18 haplotype 2 chromosome 5 centromere displayed a modular HOR architecture with defined repeat families and orientations. CpG methylation and Pore-C mapping revealed epigenetic heterogeneity and spatial clustering of repeats, respectively, while nucleotide identity analysis showed mostly homogeneous sequences interspersed with variable regions. Together, these data highlight that centromere structure and function arise from the coordinated interplay of sequence, epigenetic modifications, and 3D genome organization (Fig. 5D).

## Discussion

Telomere-to-telomere (T2T) assembly of diploid human genomes has thus far relied on the integration of multiple sequencing technologies, typically combining PacBio HiFi reads for high consensus accuracy, Oxford Nanopore ultra-long reads for repeat resolution, and chromatin conformation capture data from short-read platforms (Hi-C) for scaffolding and phasing. Here, we demonstrate that high-quality diploid T2T assemblies can be generated using ONT as a sole sequencing platform based only on UL reads and Pore-C data. This streamlined approach reduces laboratory complexity, cost, and infrastructure requirements, while achieving assembly continuity and accuracy comparable to multi-platform HPRC assemblies.

Across 23 individuals, we reconstructed 360 fully gapless chromosomes and an additional 446 near-complete T2T scaffolds using at minimum three UL and one Pore-C flow cell. As shown before for a single genome^13^, we demonstrate that Pore-C data alone is sufficient for chromosome-scale haplotype phasing. In direct comparison with trio-based phasing, Pore-C phasing produced similar switch error rates and scaffold continuity, without requiring parental sequencing or additional technologies such as Hi-C or Strand-seq on short-read platforms. This substantially simplifies diploid genome assembly workflows and enables scalable phasing in cohorts where parental material is unavailable.

A key advance of this work is the reconstruction of diploid genomes reaching a median consensus accuracy above QV 50 without using Duplex sequencing or external polishing. These results indicate that *HERRO* error-corrected R10.4 UL reads can substitute for PacBio HiFi reads in graph-based assembly frameworks such as *Verkko*. While hybrid assemblies remain the current gold standard, our results show that Nanopore-only assemblies can achieve similar gene completeness, structural integrity, and phased variant detection performance.

Despite these advances, limitations remain. Base-level accuracy of Nanopore-only assemblies, although approaching QV50, remains modestly below that achieved with HiFi-based assemblies, particularly in homopolymeric regions where short indel errors persist. While polishing strategies (e.g. APK polishing kit) may further reduce these errors, our results suggest diminishing returns relative to increased sequencing costs. Continued improvements in pore chemistry, base calling models, computational error correction models and assembly algorithms will likely further narrow this gap.

Assembly continuity was highest for metacentric and submetacentric chromosomes, whereas acrocentric chromosomes and certain centromeres remained challenging. Although complete resolution of individual acrocentric chromosomes was achieved in selected samples, repetitive ribosomal DNA arrays and large satellite blocks continue to represent the most difficult regions for automated assembly. Advances in UHMW DNA extraction, further increases in read length distribution, and improved repeat-aware assembly algorithms may ultimately resolve these remaining gaps. Indeed, we observed a steady increase in assembly quality over the course of this study, which is mainly attributed to the increase in yield of reads longer than 100 kb.

Importantly, we showed that the use of a single sequencing platform enables simultaneous reconstruction of genome sequence, haplotype phase, methylation patterns, and 3D chromatin organization. By integrating phased SNVs with Pore-C contact data, we generated haplotype-resolved maps of TADs, revealing allele-specific chromatin structures at imprinted loci and clear signatures of X-chromosome inactivation. Nanopore reads are more efficient for phased applications than Hi-C reads for two reasons: their longer fragments span more heterozygous variants, improving phasing; and individual reads capture multi-way contacts, increasing informative linkage. Together, these features improve haplotagging and reduce the sequencing required to resolve allele-specific chromatin structure at TAD-scale resolution compared with Hi-C-based approaches. This integrated framework extends beyond structural genome assembly and provides a foundation for functional diploid genomics.

The availability of phased T2T assemblies opens new opportunities for studying structural variation, centromere evolution, regulatory architecture, and population diversity. While short-read whole-genome sequencing captures approximately 93% of the human genome^25^, unresolved regions are enriched for structural variants, satellite repeats, and regulatory domains that may contribute to rare diseases and cancer predisposition. Alterations in TAD boundaries and higher-order chromatin structure have already been implicated in developmental disorders^26^. Systematic generation of haplotype-resolved T2T assemblies with chromatin topology across diverse populations may therefore provide a more complete reference framework for interpreting pathogenic variation.

At the same time, clinical translation will require further improvements in cost efficiency, turnaround time, and standardized analysis pipelines. Although our workflow reduces the need for multiple sequencing platforms, UL DNA preparation remains technically demanding and sensitive to sample quality. Continued protocol optimization and automation will be critical for broader adoption.

In summary, we present a scalable, Nanopore-only workflow for diploid T2T genome assembly that integrates sequence, methylation, and chromatin topology from four flow cells per individual. By lowering technical and infrastructural barriers, this approach facilitates expansion of T2T genomics beyond large consortia and supports population-scale and functional genomic studies. Ongoing efforts to expand this resource to 50 publicly available T2T genomes will further contribute to the development of comprehensive pangenome references and functional diploid genome maps.

## Methods

### Cohort description

Genomes of 23 healthy adults were assembled, representing diverse ancestries including European, Latin American, South Asian, and East/Southeast Asian backgrounds. The cohort comprised 11 females and 12 males (mean age 45.9 years; range 25–95 years). Several participants reported mixed ancestral origins. Two individuals (T2T00 and T2T04) were sequenced together with their parents to enable trio-based phasing. Informed consent was obtained from all patients or their guardians for use of their samples for research, as approved by the ethics committee of the University Hospital Tuebingen in accordance with to the Declaration of Helsinki (ethics approval No. 113/2025BO1).

### DNA extraction and ONT UL Sequencing

Ultra-high molecular weight (UHMW) DNA was extracted from fresh EDTA blood, collected in EDTA anticoagulant tubes, using the NEB High Molecular Weight DNA Extraction Kit for Cells & Blood (New England Biolabs, T3050) according to the manufacturer’s protocol with optimizations for ultra-long read (ULR) sequencing. The UHMW DNA was eluted in ONT Extraction Elution Buffer (EEB) and incubated at room temperature for several days to achieve complete homogenization before library preparation. For each individual, two ultra-long libraries were prepared from 30–40 μg of unsheared DNA eluted in 750 μL of EEB buffer using the ONT Ultra-Long Kit V14 (SQK-ULK114). Libraries were pooled and sequenced across three PromethION R10.4.1 flow cells (FLO-PRO114M) at a total loading volume of 150 μL per load (75 μL SBU, 7.5 μL LSU, 67.5 μL library). Flow cells were periodically washed and reloaded with fresh libraries up to 5 times when active pore counts declined below ∼25% to maximize throughput.

### Pore-C Library Preparation and Sequencing

Peripheral blood mononuclear cells (PBMCs) were isolated from fresh peripheral blood using Ficoll density-gradient centrifugation and processed following the Pore-C protocol. Briefly, 20 million cells were crosslinked with 1% formaldehyde for 10 minutes, digested overnight with restriction enzyme NlaIII, and proximity ligated using T4 DNA ligase for 6 hours. Subsequently, proteins were digested overnight with proteinase. Ligated genomic DNA was isolated and purified using self-prepared SPRI beads. After reverse crosslinking and purification, 2.4 μg DNA was prepared using the ONT Ligation Sequencing Kit V14 (SQK-LSK114 XL) with modifications to the manufacturer’s protocol aimed at enhancing efficiency. The DNA was further processed in a series of steps, including nick repair and end preparation, to facilitate the attachment of sequencing adapters. Pore-C libraries were sequenced on PromethION R10.4.1 flow cells (typically 1-2 per sample; five for selected samples). Flow cells underwent multiple nuclease washes and reload cycles.

### Basecalling, Error Correction, and Read Filtering

All raw data were basecalled using *Dorado v1.0.2* with the R10.4.1 super-accuracy (SUP) model (dna_r10.4.1_e8.2_400bps_sup@v5.2.0). 5mC and 5hmC modifications were called using 5mCG_5hmCG@v1. Ultra-long reads were error corrected using *dorado correct* (*herro-v1* model). These corrected reads were used as high-quality (HQ) input reads in place of PacBio HiFi reads for assembly. Read statistics were computed using *FastCAT v0.15.2* and *dorado summary*. Reads were filtered and subsampled using *filtlong* to 50x coverage for HQ reads and 70× for UL reads. Minimum read length thresholds were 10 kb (HQ) and 80 kb (UL). A minimum read quality score of 9 was applied. We used the parameter *--length_weight 10* for *filtlong* to favor length over quality in read subsampling.

For genome assembly, we used a single Pore-C flow cell for which we copied the FASTQ files to the assembly input folder. For *Verkko* assembly this file was used without preprocessing, while a pseudo-paired-end Hi-C format was generated for the *hifiasm* assembler using a custom python script. To this end, Pore-C reads were split at CATG motifs and the resulting fragments combined with up to 5 other fragments of the same read into a set of pseudo-paired-end reads.

### Genome assembly Verkko Assembly

Genome assemblies were generated using *Verkko v2.2.1*^27^ in local mode on a single compute node per sample. Corrected HQ reads were supplied as *--hifi* input and ultra-long reads as *--ont*, with correction disabled (*--no-correction*). To compare *Verkko* results with different phasing options, assemblies were generated in: (1) Unphased mode; (2) Pore-C phasing mode (--*porec*); (3) Trio-aware mode using haplotype-specific k-mers generated with *Meryl* and *Merqury*.

For trio binning with *Verkko*, we first counted homopolymer-compressed k-mers (k=30) for each parent and child using *meryl count compress*. Then, we extract haplotype-specific k-mers for each parent using *hapmers.sh* from the *Merqury* package. These parental k-mer sets were used as input for *Verkko’s--hap-kmers* option enabling trio-aware scaffolding and phasing.

### Hifiasm Assembly

For samples T2T00 and T2T04, trio-based and Pore-C-based assemblies were generated using *hifiasm*^9^ with ONT error correction enabled *(--ont*) and telomere motif detection (*--telo-m CCCTTA*). *Hifiasm* assembly with Pore-C phasing was performed using the parameters: *--telo-m CCCTTA --dual-scaf --ul-cut 70000-l2 --ont --h1 {porec_r1}-h2 {porec_r2} {ul}*.

For *hifiasm* trio binning of the samples T2T00 and T2T04, we generated k-mer databases (k=37) for each parent using *yak count*. The resulting .yak files were supplied to *hifiasm* with the −1 and −2 options for maternal and paternal k-mers. Hifiasm with trio binning was performed using the parameters: *--telo-m CCCTTA --dual-scaf --ul-cut 70000 −l0 --trio-dual --ont −1 {maternal} −2 {paternal} {ul}*.

### T2T Classification

Classification of chromosome sequences as T2T-contig or T2T-scaffold is determined by the presence of telomeric motifs, full alignment to a reference chromosome and the presence of gaps (stretches of NNN sequence). We classified chromosomes as T2T-Scaffolds if: (1) Telomeric repeats were present at both ends; (2) Alignment to T2T-CHM13 covered ≥95% of a chromosome in a single block. If T2T-Scaffolds were also gap free (no N-stretches) they were classified as T2T-Contigs.

To compute T2T-CHM13 coverage we created a pairwise alignment in PAF format using *minimap2* and then used *minigraph cov_cal* to identify sequences overlapping >=95%. A custom python script was used to identify and count N-stretches in the assembled sequences and determine the final classification. The script provides summary tables showing the number, size, and location of gaps.

### Assembly quality control

#### Assembly Integrity

Potential mis-assemblies were identified using *NucFlag* and *Flagger*, based on HQ read re-alignments. Error corrected HQ reads were re-aligned to the assembled genomes using *minimap2* with the parameter preset *lr:hqae*. Mapped reads were analyzed with *NucFlag* using default options. Plots were created for all regions using the Repeatmasker annotation. *Flagger* was executed from a docker container in a two-step process. First, *flagger bam2cov* was used to create a coverage file from the re-alignment. Second, *hmm_flagger* was used to generate a bed file with potential integrity issues.

#### Genome completeness

We assessed the completeness and quality of our T2T assemblies by comparing against the T2T-CHM13 reference genome^1^. Ensembl release 113 cDNA sequences were aligned using *minimap2* with options *-cxsplice* and *-C5* to both the assembled genome and to T2T-CHM13. The number of partial, complete and duplicate occurrences of every transcript were assessed using *paftools asmgene*. Genome completeness was then calculated by dividing the sum of complete and duplicate alignments in the asmgene output by the sum of full alignments in the reference. Furthermore, the numbers of missing multi copy genes (MMC) and missing single copy genes (MSC) were calculated using *paftools asmgene*. MMC is calculated by finding the number of multi copy genes in the assembly divided by the number of multi copy genes found in the reference. MSC is the fraction of single copy genes found in the assembly to the number of single copy genes found in the reference.

#### Consensus quality

To determine QV values we first performed short read WGS (srWGS) for all samples using Illumina NovaSeq X+. QV values were then generated using *Merqury* based on srWGS re-alignments to the assembled genomes. QV values were calculated for three kmer lengths (k=21, k=31, k=37) for each generated assembly.

### Variant calling and phasing accuracy

UL reads were aligned to T2T-CHM13 v2^1^ using minimap2^28^. Sex was inferred from X/Y coverage ratios (>0.5 results in male). Coverage profiles were calculated using *mosdepth*^29^. Fully phased single nucleotide variants and indels were called with *dipcall*^30^, larger structural variants with *hapdiff*^31^. Phasing accuracy was evaluated using *whatshap compare* against the GIAB CHM13 phased benchmark (CHM13v2.0_HG2-T2TQ100-V1.1^2^). We used *whatshap stats*^32^ to generate variant statistics. Further variant statistics were extracted from VCF files generated by *hapdiff* and *dipcall* using custom python scripts.

### Ancestry analysis

Variants were normalized using *bcftools norm* and intersected with a lifted 1000 Genomes panel in T2T-CHM13 coordinates using *bcftools view-T*, as suggested by Lalli et al.^33^. Normalized VCFs were filtered to retain only biallelic SNVs and indels, excluding spanning deletions and large variants exceeding 50 bp in either reference or alternate allele length. Sample and reference VCFs were merged using *bcftools merge* with options *--merge* none to prevent forced merging of incompatible variants, and *--missing-to-ref* to impute missing genotypes as homozygous reference calls. The merged VCF was converted to PLINK2 binary format using *plink2 --vcf*. Chromosome codes were standardized with *--split-par b38* to handle pseudoautosomal regions on chromosome X, and half-calls were set to missing (*--vcf-half-call m*). Variant IDs were standardized using the format chr:pos (*--set-missing-var-ids @:#*). Quality control filters were applied to the merged dataset using plink2 with the following lenient thresholds optimized for ancestry analysis: minor allele frequency (MAF) ≥ 0.01 to retain rare variants informative for population structure, per-variant missingness ≤ 15% (*--geno 0.15*), and Hardy-Weinberg equilibrium p-value ≥ 0.001 (*--hwe 1e-3*), with the latter threshold intentionally lenient to accommodate population structure and potential familial relationships. Variant IDs were updated to a standardized format (chr:pos:ref:alt) to ensure consistency across the dataset.

PCA was performed using PLINK2 following LD pruning (r²=0.2). Ancestry inference was performed using *RFMix*^34^. Variants in strong linkage disequilibrium were removed using *plink2 –indep-pairwise* with a 50-SNP sliding window, 10-SNP step size, and r² threshold of 0.2. PCA was performed using *plink2 –pca* to extract the top 20 principal components. The resulting eigenvectors and eigenvalues were used to visualize population structure, stratified by either population or super-population labels of 1000GP. We used *RFMix* to identify local and global ancestry of the subjects with parameters *crf-spacing=0.001* and *rf-window-size=0.1*.

### Haplotype-phased 3D genomes using Pore-C

Pore-C data were processed using *wf-pore-c*^35^ and *pairtools*^36^ to generate contact maps (pairs files). Haplotype tagging of Pore-C reads was performed as described by Chen et al.^19^. In brief, reads were mapped to the reference genome using *Falign*. SNVs and indels in high quality regions with >=5x Pore-C coverage were used for *happlotagging* with *dip3d haplotag*. Inter-chromosomal contacts were not included in the phasing. Finally, separate alignment files for each haplotype were extracted. Haplotype-separated Pore-C alignment files were processed into pairs format using *pairtools parse2*^36^. To count correct non-adjacent contacts on the same fragment we used the *--expand* and *--flip* options. Contact maps from multiple flow cells of the same sample were merged with *pairtools merge*. Statistics were calculated with *dip3d stats*. Juicebox^37^ was used for visualization of contact maps.

For further analysis we converted the pairs files to *Cooler*^38^ format using *cooler cload pairs* with a minimum binwidth of 1.000 and then created a mcool file using *cooler zomify* with aggregate resolutions for binwidths 1.000, 5.000, 10.000, 50.000, 100.000 bp. Finally, we created a corrected matrix using the ICE algorithm with cooler balance that serves as basis for detection of 3D features. We used *cooltools insulation* to calculate insulation scores for the resolutions 25000bp and 40000bp and *cooltools eigs-cis* to create a compartment track. For TAD and Loop calling we used *hicExplorer* tools^39^ again starting from the balanced pairs file. Then we used *hicFindTADs* to create a TAD score and Boundary plot for the resolutions.

### Haplotype-resolved Methylation Analysis

Modification-aware UL reads were subsampled to a maximum size of 100 Gigabytes and aligned to both the assembled genomes and the T2T-CHM13 reference genome with minimap2. Reads were haplotagged with phased SNP calls using *longphase haplotag*. Reads that could not be confidently assigned remained untagged (HP:0). Haplotagged reads were split into separate BAM files for each haplotype using *samtools view -d HP*. Haplotype-specific methylation frequencies were computed with *modkit* and exported as bigWig files. In brief, case modification consensus was generated using *modkit pileup*, summary statistics were generated by *modkit summary*, and bedMethyl files were converted to bigWig format using *modkit bedmethyl tobigwig*, enabling visualization of CpG methylation in genome browsers on both T2T-CHM13 and T2T-sample coordinates.

### Centromere and telomere structures

Centromeric regions were characterized using *CenMap* tools with default settings on all 23 samples as described in Logsdon et al.^4^. *CenMap* finds satellite repeats, annotates the High Order Repeat (HOR) classification, identifies centromeric dip regions (CDRs), and assesses assembly correctness of the centromeres using *NucFlag*. We visualized the HOR structure using *cenplot* in a custom configuration.

## Data availability

Raw sequencing data, genome assemblies, and the generated annotations in this study are available at https://github.com/imgag/T2T_Genomes. All the code used for the analyses is available at https://github.com/imgag/T2T_ONT.

## Acknowledgements

This work was supported by grants via the BEGIN program of Baden-Württemberg to OR (MWK25-04-3214/2/8) and by the European Commission (GoE: 101168231) and the BMFTR (GoE: 01KX2526A) via the “Genomes of Europe” initiative to OR. Part of the work was supported by the German Research Foundation (DFG) to FC (CH2339/5-1). We thank Ms. Marion Loitz for assistance in blood sampling and all T2T participants for their blood donation. We kindly acknowledge the technical contributions of Yogesh Singh and Wenxu Zheng during the Pore-C implementation phase, Michaela Pogoda for assistance with Illumina sequencing, and Jakob Admard for making all the datasets available online. Furthermore, we thank Alexander Vogel from ONT for helpful discussions throughout the course of the project, as well as the members of the HPRC for stimulating scientific exchange.

## Authors contributions

O.R., S.O., B.K., and C.G. designed and led the study. R.P., F.C., H.S.H., E.B.A. performed sample extraction, library prep and sequencing. C.G. and S.L. implemented the computational analysis pipeline and performed data analyses. N.C. carried out resource management. C.G., S.O., O.R. wrote the main manuscript, with revisions by all authors. All authors have reviewed and approved the final manuscript.

## Use of AI and language tools

All authors confirm that this manuscript was conceived, prepared, and written by the authors. The content reflects the original work, ideas, and analyses of the authors. A professional language editing tool (DeepL) and OpenAI was used for grammar and sentence structure corrections.

## List of Abbreviations

3C: Chromatin conformation capture
αSAT: alpha satellite
bp: base pairs
CDR: Centromeric Dip Regions
DNA: Deoxyribonucleic Acid
GHGA: German Human Genome Archive
GIAB: Genome in a Bottle
HOR: high-order repeat
HQ: High Quality
HPRC: Human Pan-Genome Reference Consortium
kb: kilobases
MMC: missing multi copy gene
MSC: missing single copy gene
ONT: Oxford Nanopore Technologies
QC: Quality Control
SNV: Single Nucleotide Variant
srGS: short read WGS
SV: Structural Variants
T2T: Telomere-to-Telomere
TAD: Topologically Associating Domain
UHMW: Ultra high molecular weight
UL: Ultra Long Nanopore Reads
WGS: whole genome sequencing
QV: Quality values

