## Supplementary material for "Single-Platform Nanopore Sequencing Enables Diploid Telomere-to-Telomere Genome Assembly and Haplotype-Resolved 3D Chromatin Maps": Gross et al 2026_Supplement.pdf

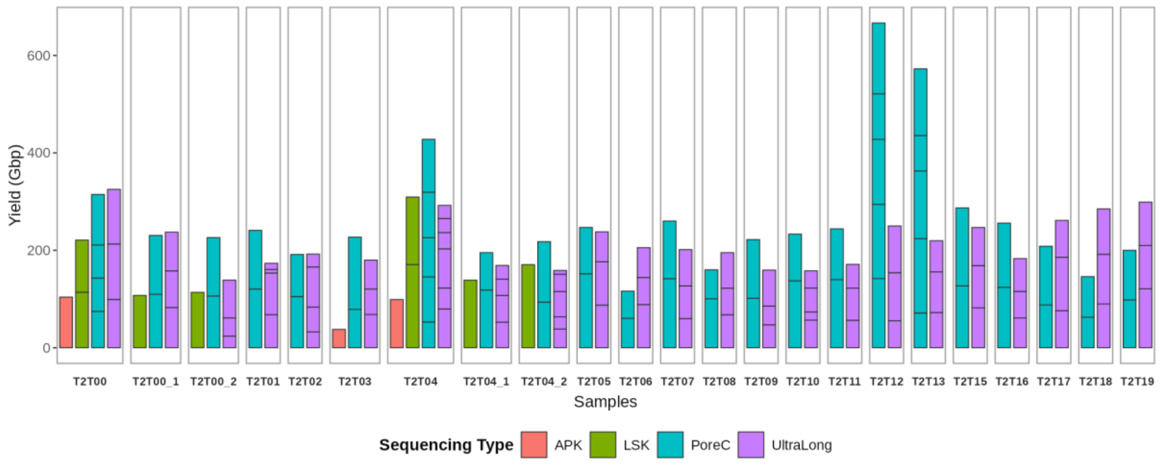


***Supplementary Figure 1: Sequencing yield across all ONT flow cells.*** *Overview of total sequencing output per flow cell for all samples. The standard workflow included three ultra-long (UL) and one Pore-C flow cell per individual. Additional UL flow cells were sequenced for selected samples to compensate for low-yield runs. Samples T2T00, T2T04, T2T12, and T2T13 received additional Pore-C sequencing. For trio parents, one standard-length ligation sequencing (LSK) flow cell was added to supplement the HQ dataset.*

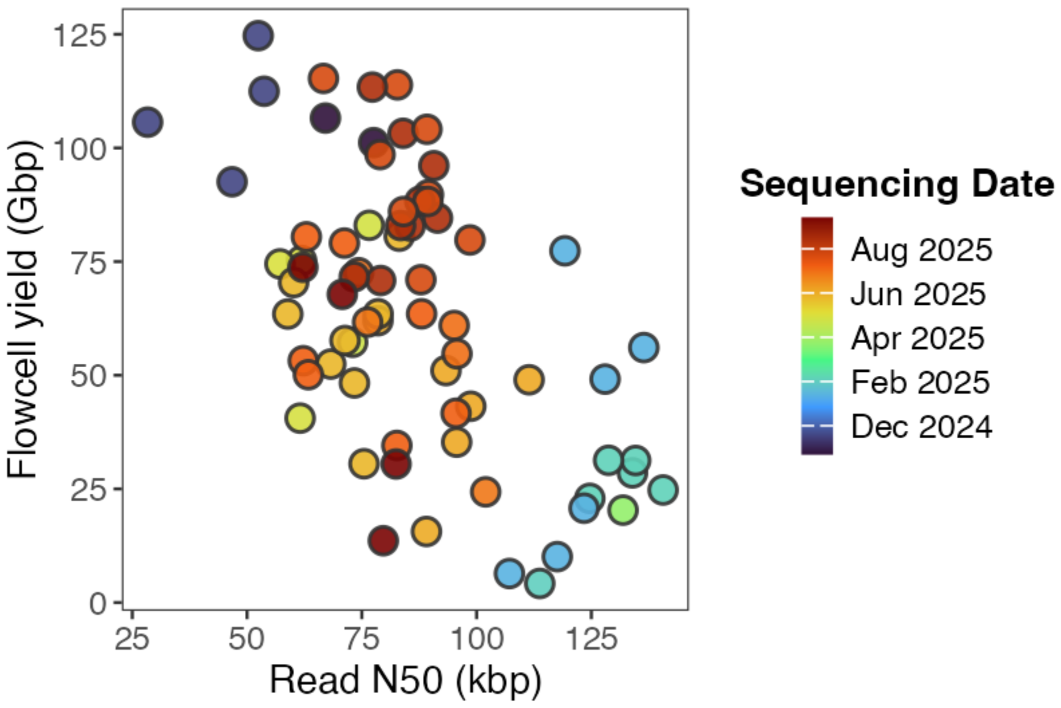


***Supplementary Figure 2: Optimization of UL read length distribution.*** *Read length distributions of UL flow cells over the course of the project. Color scale reflects sequencing date. Progressive protocol optimization improved the balance between total yield and read length N50, increasing the proportion of reads >100 kb in later samples.*

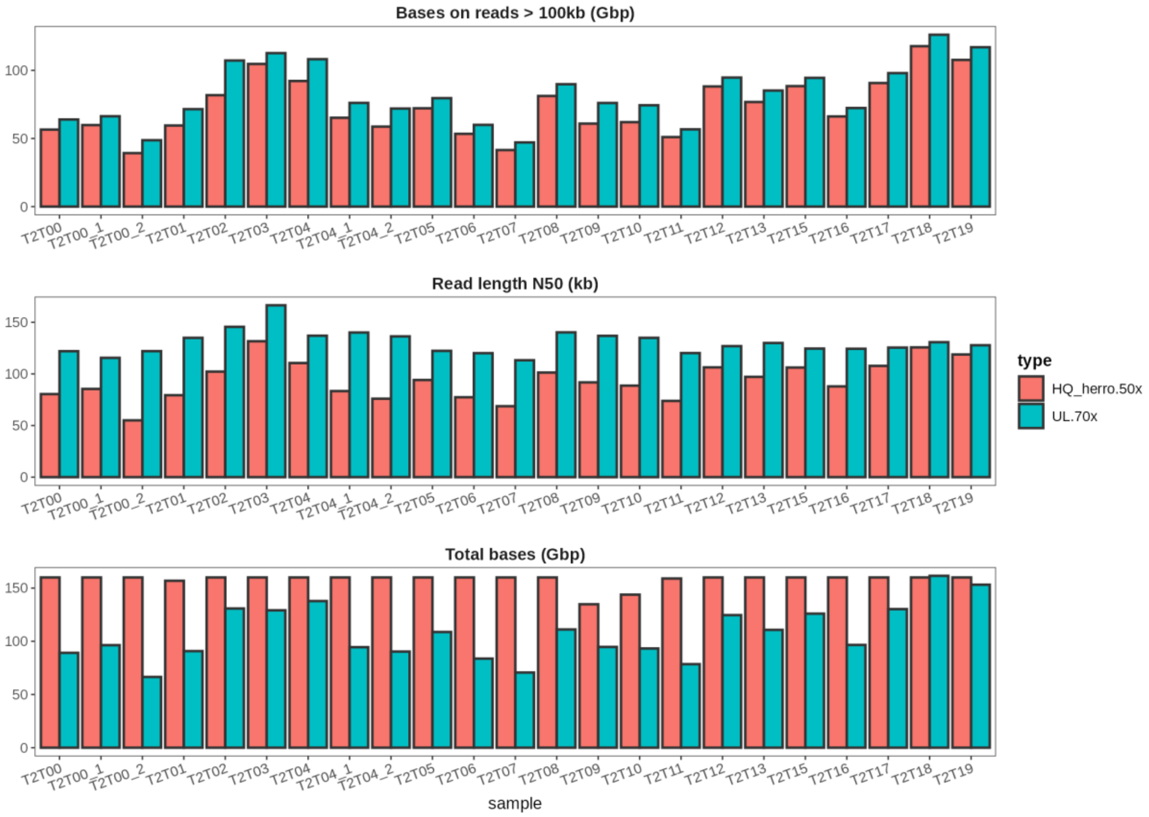


***Supplementary Figure 3. Assembly input read statistics.*** *Read statistics for combined HQ and UL datasets used for assembly. HQ reads were capped at 50× coverage; UL reads were capped at 70× coverage and filtered to retain only reads ≥80 kb. Length-based subsampling prioritized longer reads to maximize repeat resolution.*

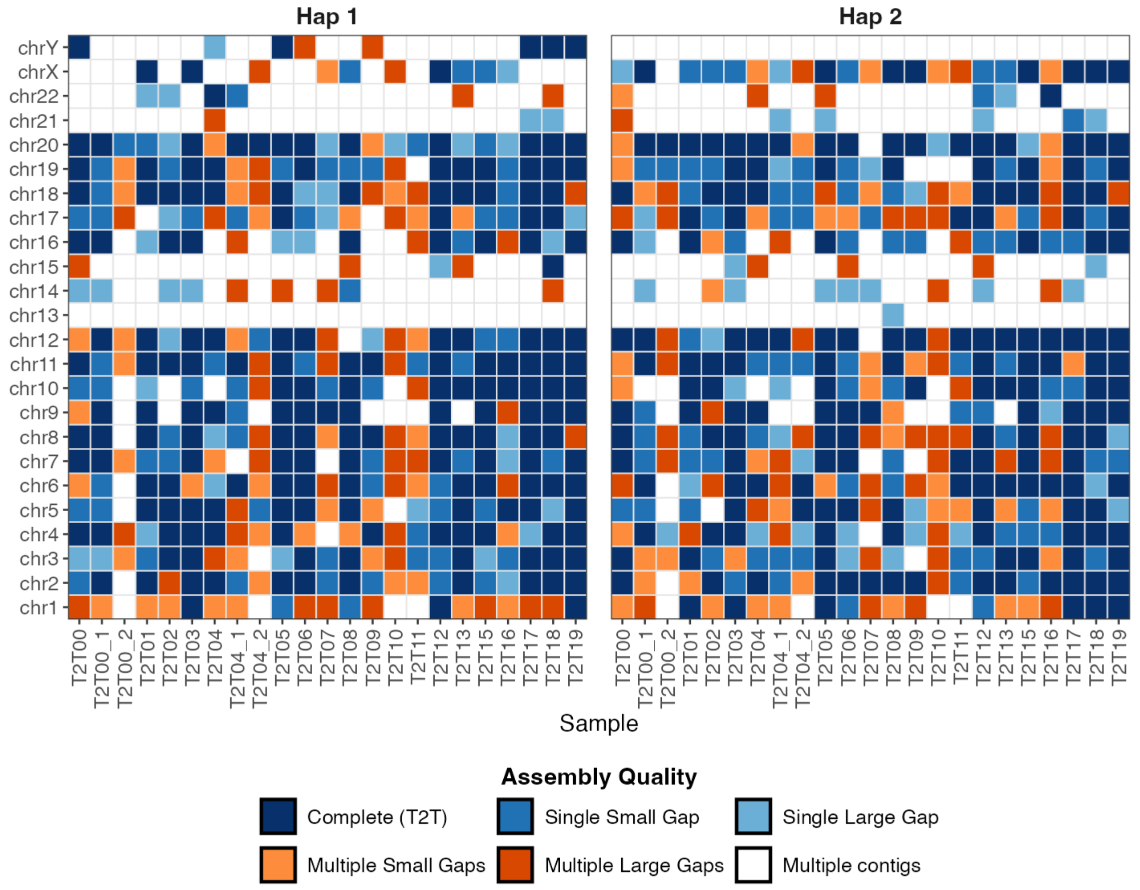


***Supplementary Figure 4. Detailed chromosome assembly classification.*** *Chromosome-level assembly status for all haplotypes with expanded categorization. Dark blue indicates gapless T2T contigs spanning entire chromosomes. Light-colored categories represent T2T scaffolds containing gaps. Large gaps are defined as single gaps >50 kb or cumulative gap size >100 kb per chromosome.*

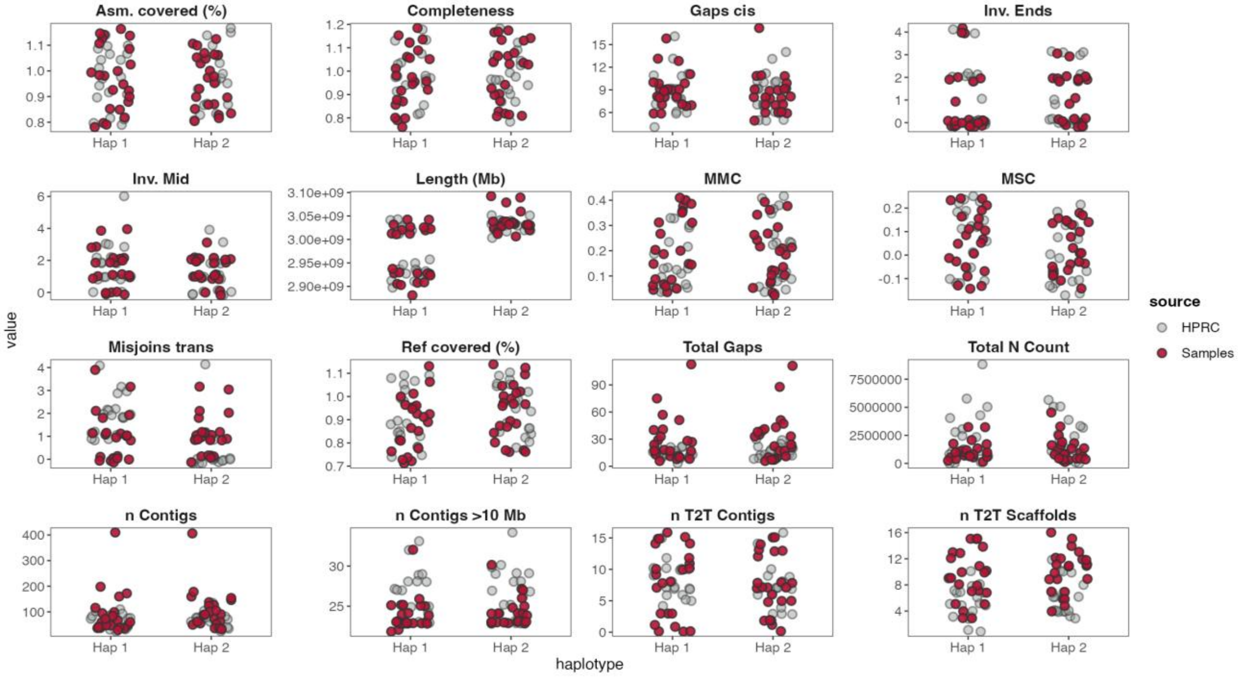


***Supplementary Figure 5. Extended reference-free assembly quality metrics.*** *Comprehensive comparison of Nanopore-only assemblies (red) and randomly selected HPRC assemblies (gray; HG002 excluded). Metrics include T2T contig and scaffold counts, gap statistics, inter- and intra-chromosomal misjoins (paftools misjoin), genome completeness, missing multi-copy genes (MMC), missing single-copy genes (MSC) (paftools asmgene), and reciprocal genome coverage statistics (paftools asmstat) relative to T2T-CHM13 v2.*

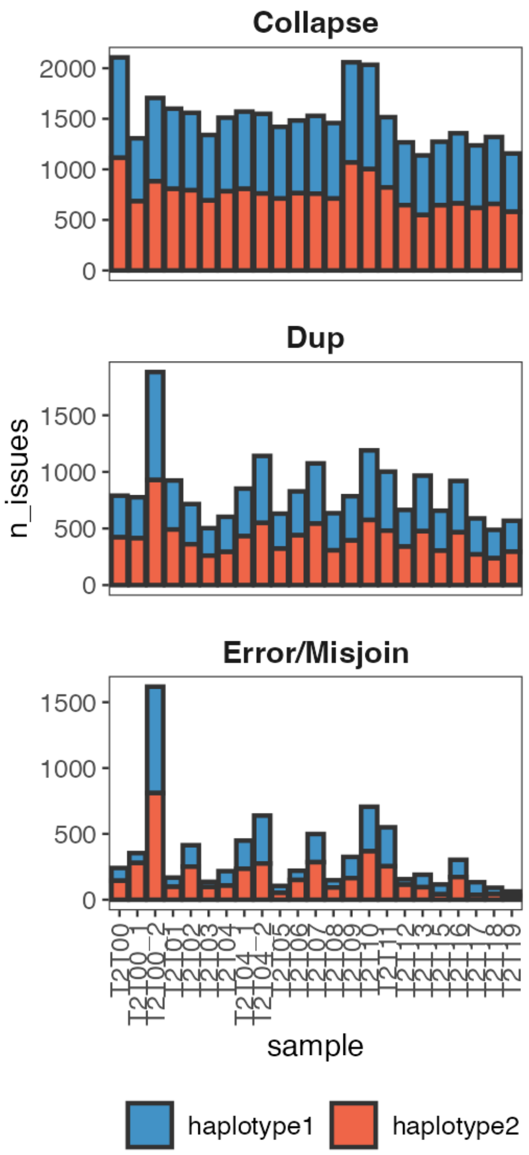


***Supplementary Figure 6. Flagger-based detection of potential assembly issues.*** *Total number of potential assembly issues identified per haplotype using Flagger based on read realignment coverage patterns.*

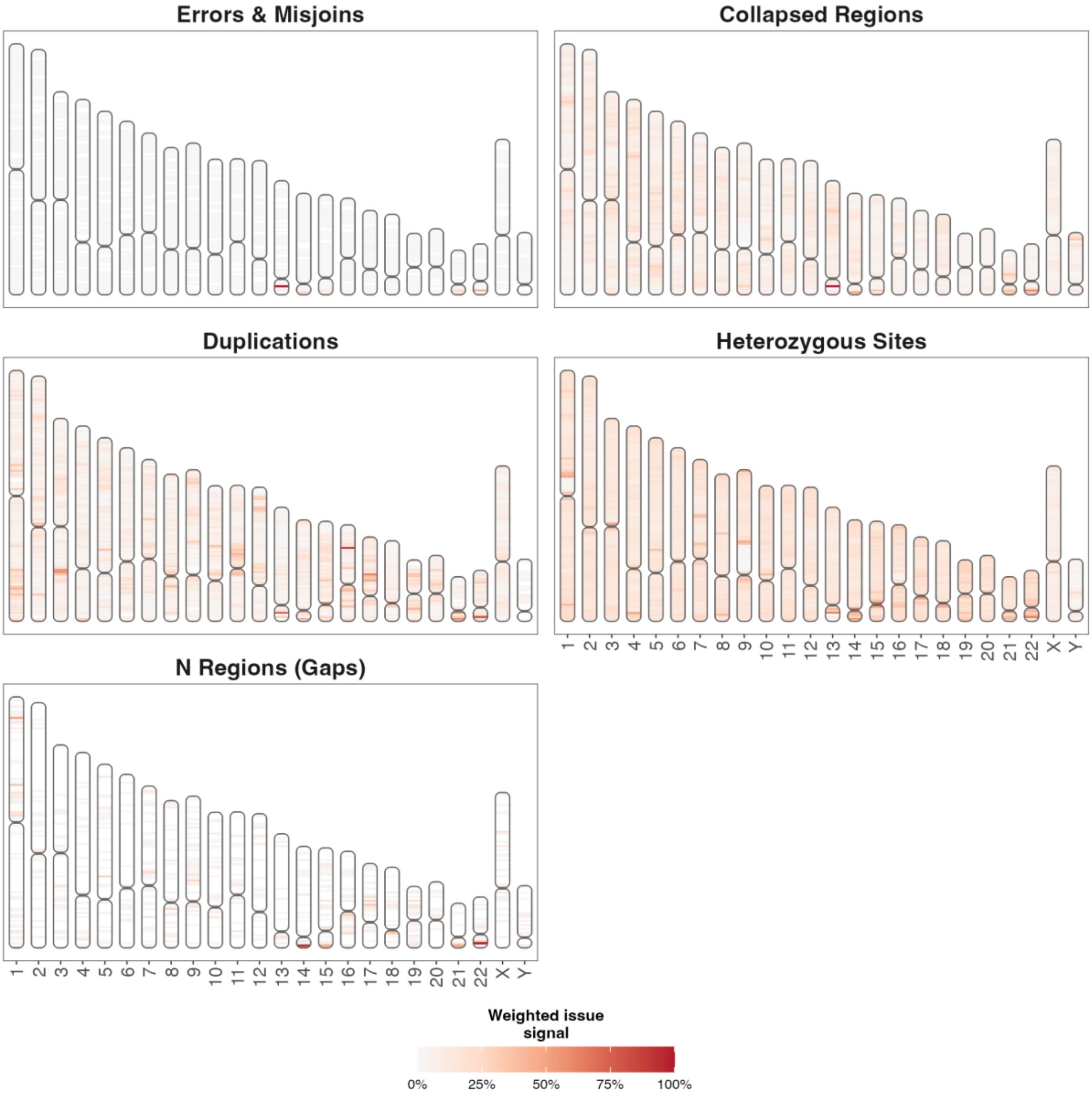


***Supplementary Figure 7. Genomic distribution of potential assembly issues.*** *Combined localization of candidate assembly issues detected by Flagger and NucFlag. Coordinates were lifted to T2T-CHM13 v2 reference positions. To avoid overrepresentation by low-quality samples, counts were normalized across assemblies.*

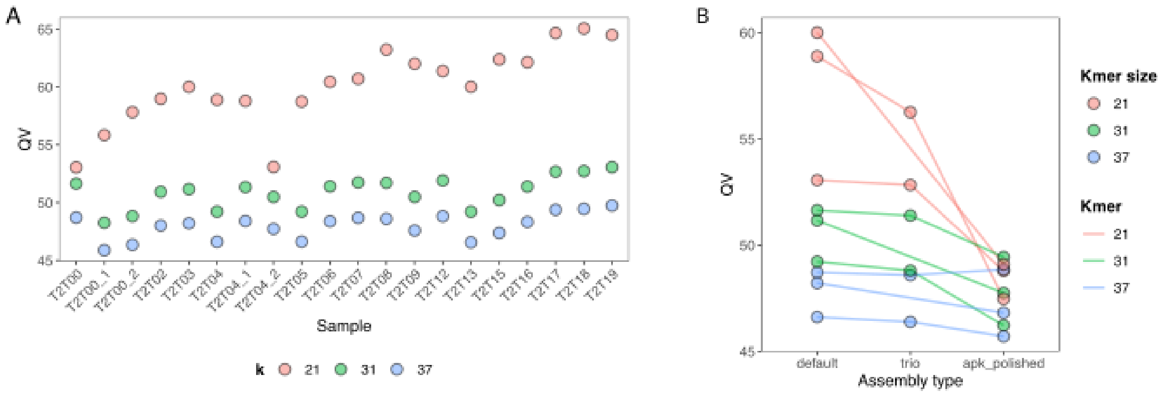


***Supplementary Figure 8. Consensus quality values (QV) across assembly strategies. (A)*** *Merqury-derived QV values calculated using Illumina NovaSeq X Plus reads for k-mer sizes 21, 31, and 37 across all assemblies.* ***(B)*** *Comparison of QV values for different assembly strategies: Verkko with Pore-C phasing (default), Verkko with trio phasing, and Verkko with Pore-C followed by APK polishing.*

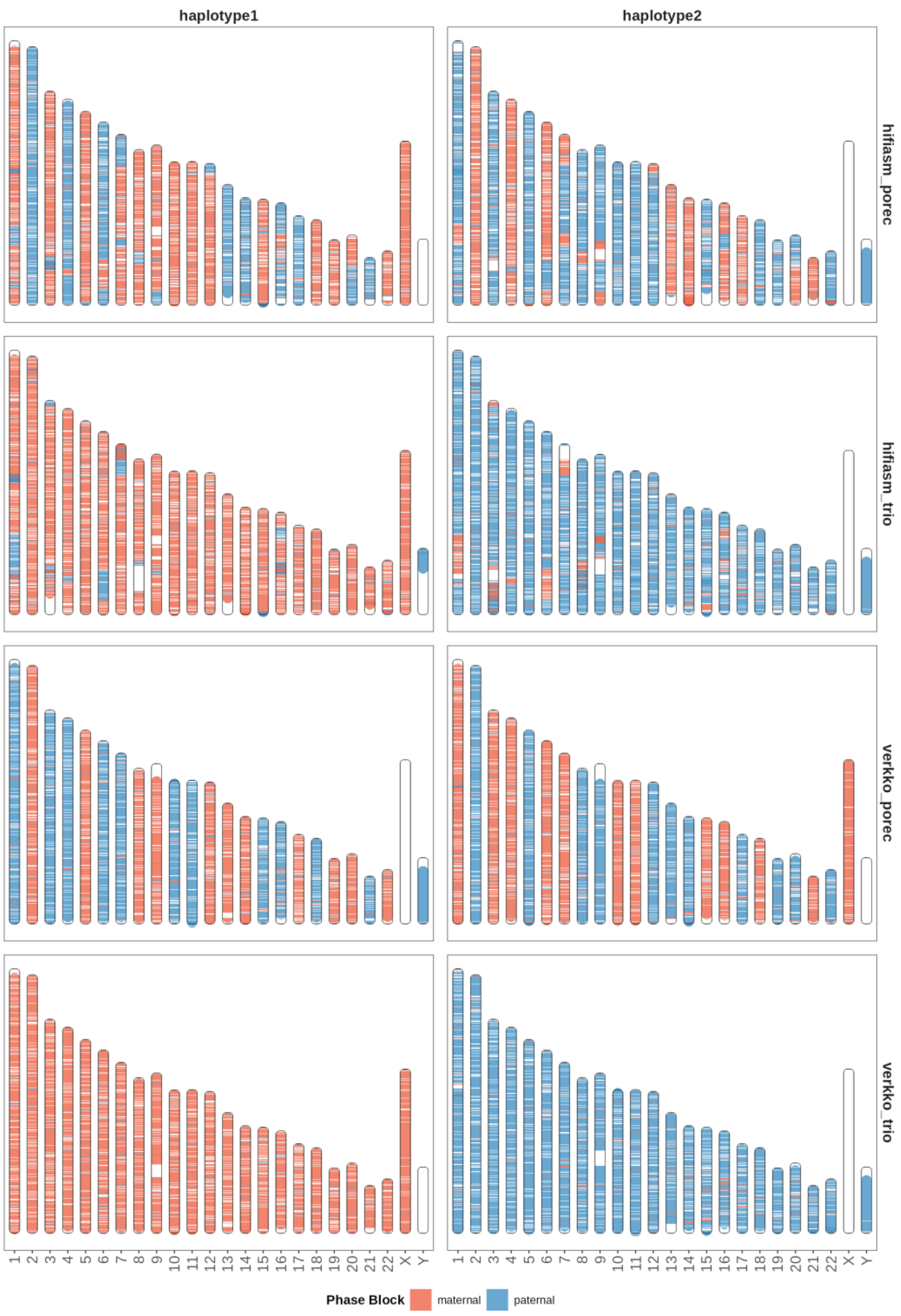


***Supplementary Figure 9. Phase block structure across assembly methods.*** *Phase block lengths for sample assemblies generated using different phasing strategies. Validation was performed using yak trioeval with identical parental k-mer sets. Pore-C–phased assemblies show chromosome-scale consistency with arbitrary parental assignment, whereas hifiasm assemblies exhibit increased large-scale phase switches relative to Verkko.*

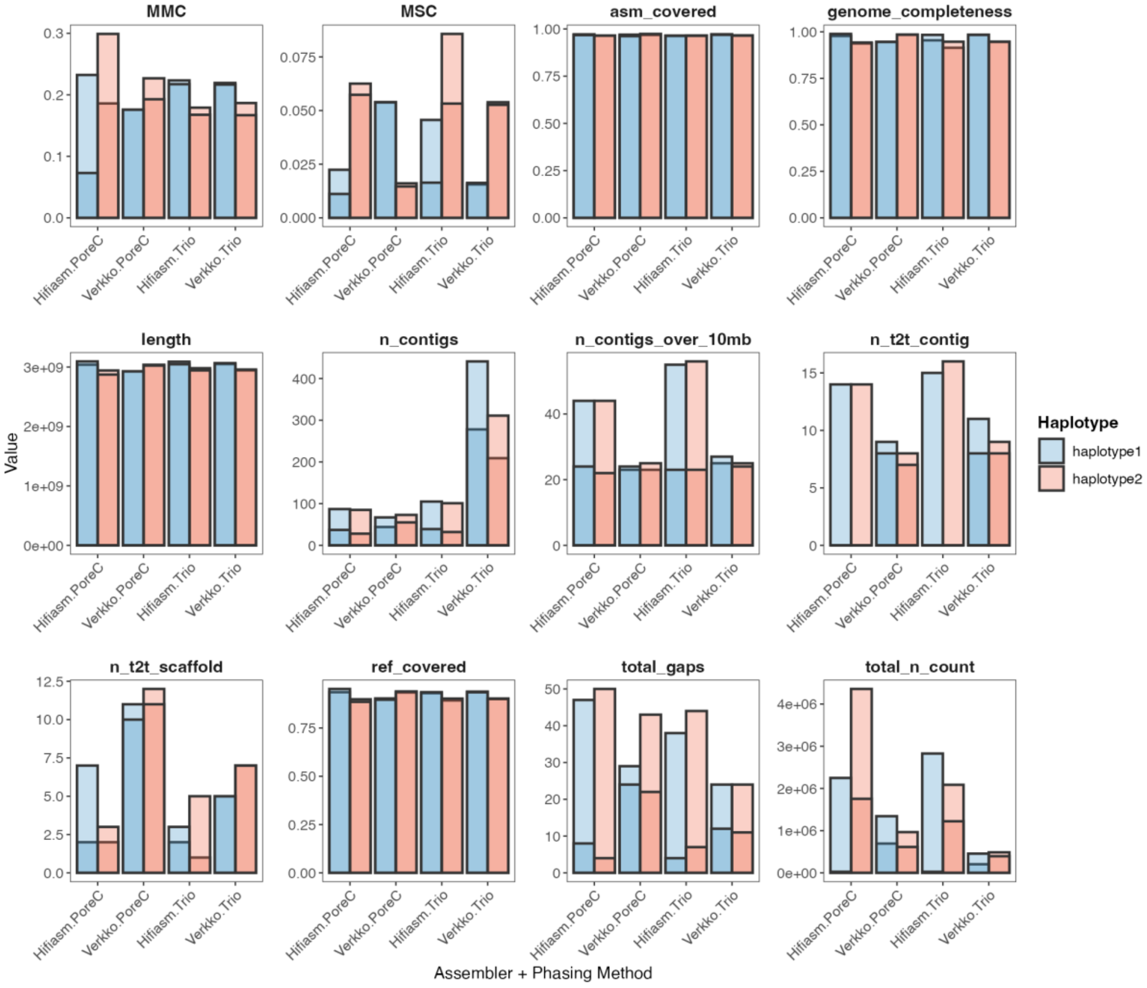


***Supplementary Figure 10. QV comparison across assembly and phasing strategies.*** *Consensus QV values for samples T2T00 and T2T04 generated using four assembly configurations: Verkko with Pore-C phasing, Verkko with trio phasing, hifiasm with Pore-C phasing, and hifiasm with trio phasing.*

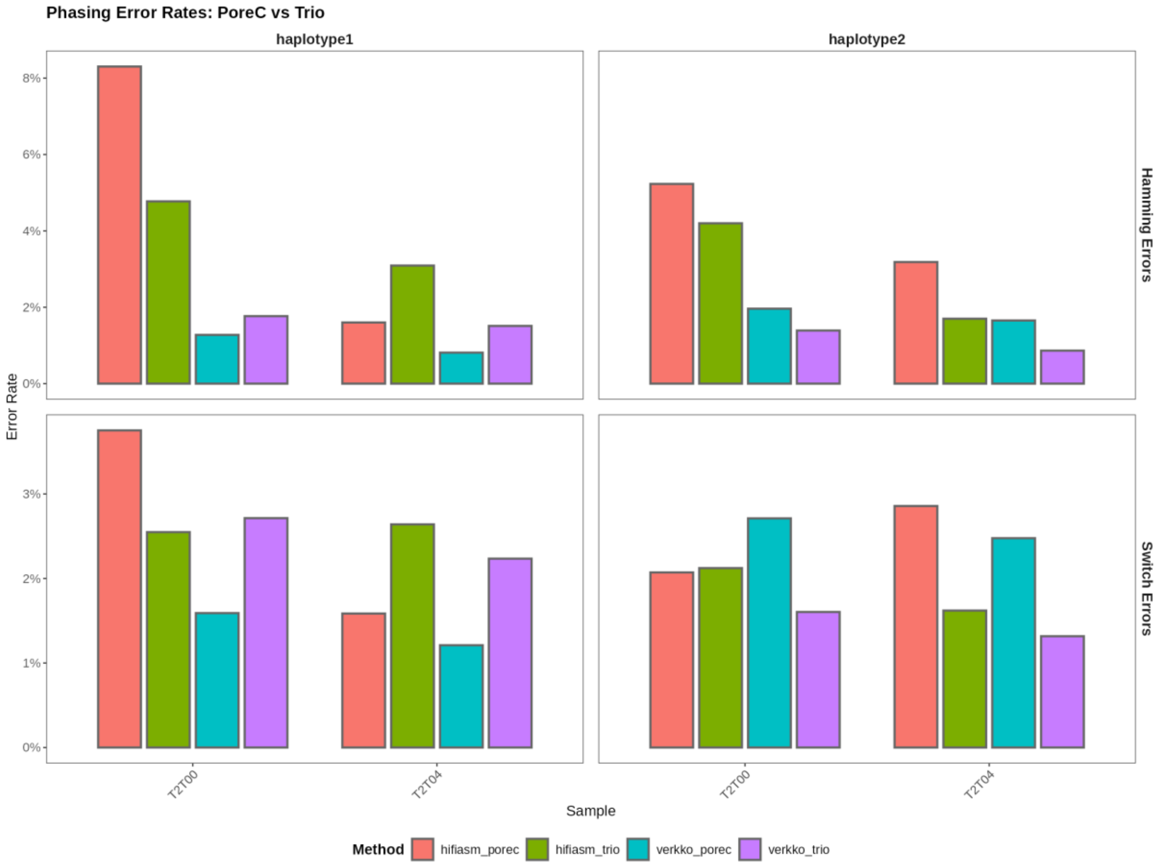


***Supplementary Figure 11. Switch and Hamming error rates across phasing methods.*** *Bar plots of Hamming error rates and switch error rates calculated using whatshap compare for Verkko (trio and Pore-C phasing) and hifiasm (trio and Pore-C phasing). Error rates are shown for both haplotypes in samples T2T00 and T2T04. Verkko demonstrates consistently lower error rates, with comparable performance between Pore-C and trio phasing.*

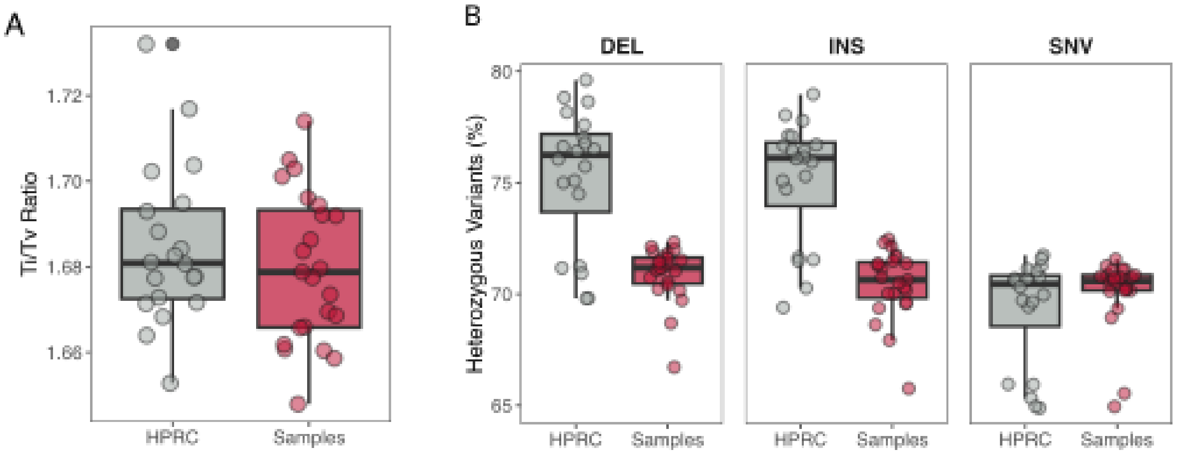


***Supplementary Figure 12. Additional small variant characteristics.*** *Comparison of variant metrics between Nanopore-only assemblies and HPRC assemblies.* ***(A)*** *Transition-to-transversion (Ti/Tv) ratios.* ***(B)*** *Proportion of heterozygous variants per genome.*

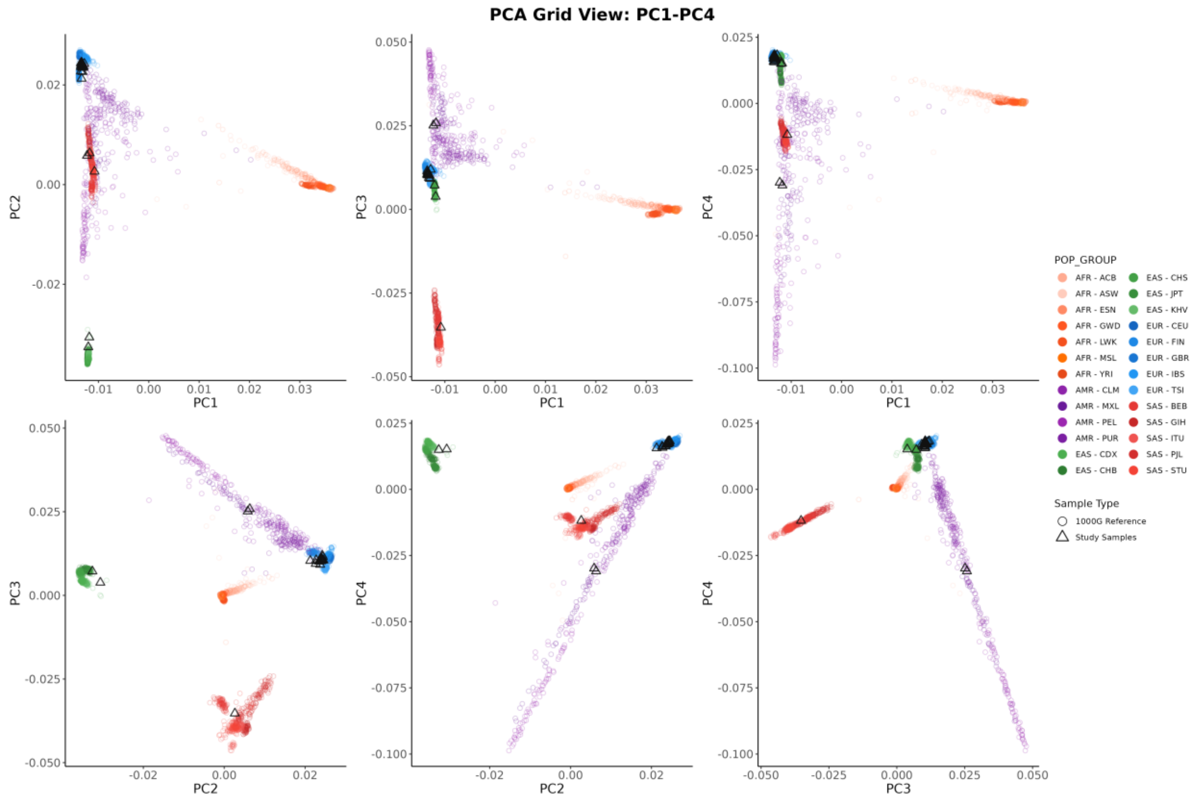


***Supplementary Figure 13. Principal component analysis of study cohort.*** *Principal component analysis of phased small variants merged with the T2T-lifted 1000 Genomes reference panel. Plots show pairwise combinations of the first four principal components. Colors represent superpopulation groups.*

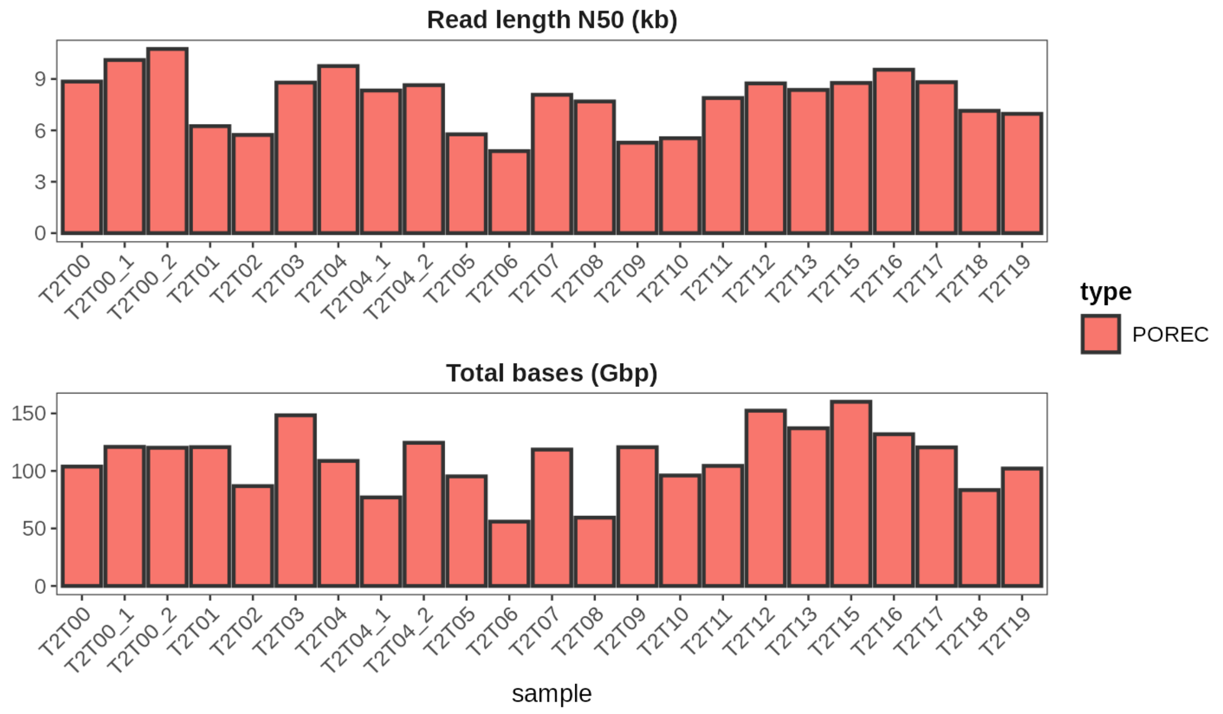


***Supplementary Figure 14. Pore-C sequencing statistics.*** *Read statistics for Pore-C datasets used for phasing and scaffolding. For assembly, a single Pore-C flow cell per sample was used. Metrics include total bases, read length distribution, and contact yield.*

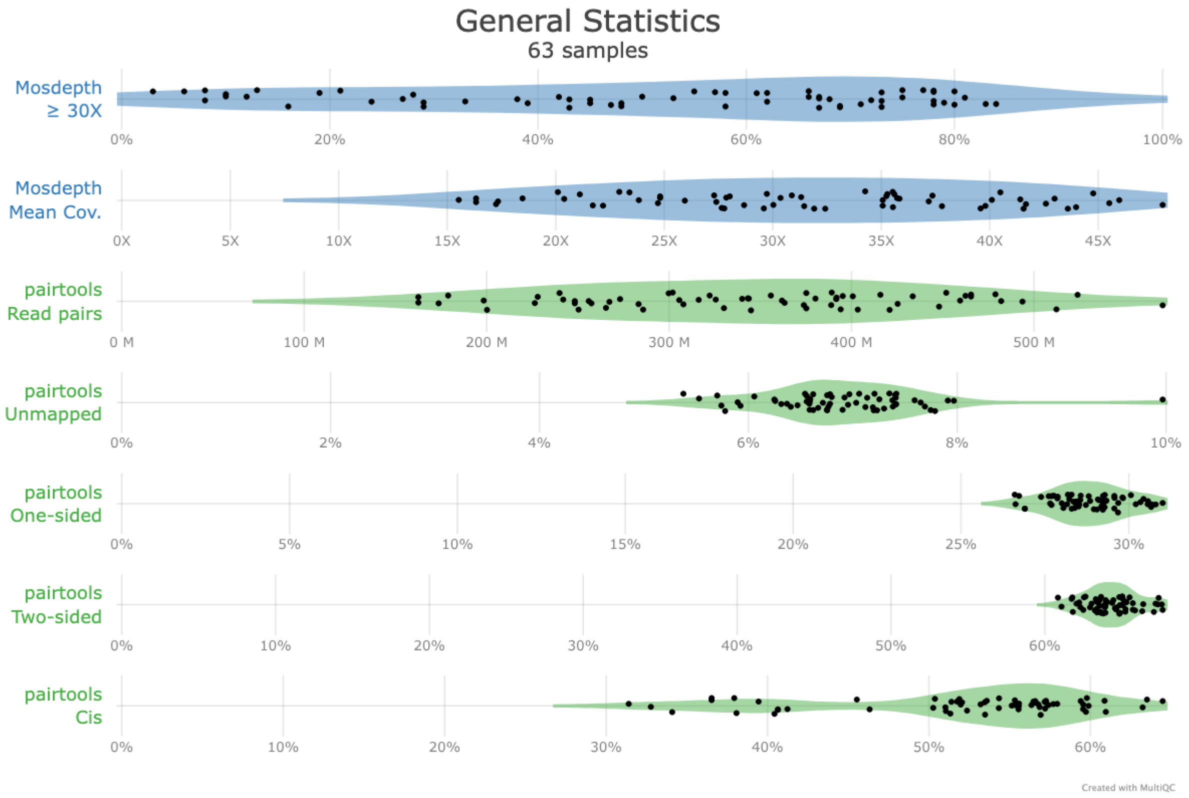


***Supplementary Figure 15. Quality metrics for individual Pore-C flow cells.*** *Quality control metrics generated by the wf-pore-c pipeline for each Pore-C flow cell. Only reads containing cis contacts were retained for haplotype phasing and downstream 3D genome analysis.*
